# Sexual Dimorphism in c-Fos Networks Governing Aggression

**DOI:** 10.1101/2025.10.06.680658

**Authors:** Antonio V. Aubry, Long Li, Romain Durand-de Cuttoli, Elizabeth Karpman, Hyoseon Oh, Arman A. Tavallaei, Scott J. Russo

**Author notes:** Equal Contribution (Long Li, Romain Durand-de Cuttoli).

## Abstract

Circuit based studies of aggression often focus on the activity of a small group of regions referred to as the “core aggression circuit,” yet a whole-brain map of activity has yet to be produced. Using resident–intruder assays in male and female Swiss Webster mice, we combined iDISCO+ c-Fos imaging with a weighted co-expression network analyses to identify mesoscale co-activation modules in aggressive (AGG) and non-aggressive (NON) animals. We performed a module preservation analysis between each phenotype within each sex followed by differential correlation to localize edge-level changes. Behaviorally, AGG mice spent more time attacking, while females showed greater social investigation. When comparing activity changes in individual regions, male AGGs showed broad activation throughout the anteroposterior axis, while females preferentially activated anterior cortical regions. Network analyses revealed that NON networks often preserved density of AGG modules, but connectivity reorganized with aggression. In male AGGs there was large spread reorganization of numerous modules: a large sensorimotor–subcortical “blue” module, a pallidal/hypothalamic/brainstem “yellow” module, and a brainstem-heavy “green” module. Surprisingly the “brown” module, enriched for classic social-behavior regions was moderately preserved in male NONs. In females, the “turquoise” module spanning somatosensory/interoceptive cortex and midbrain regions and the “red” module spanning many anterior cortical and thalamic regions were the least preserved. These results indicate that aggression recruits distributed mesoscale communities via edge-specific gain, with males displaying broad strengthening throughout the brain and females showing more targeted potentiation within anterior cortical modules. This framework nominates candidate hubs and edges for causal manipulation.

## Introduction

Aggression is an evolutionarily conserved behavior that controls social hierarchies and protects valuable resources (Lischinsky & Lin, 2020; Wei et al., 2021). In the wild, aggression is a necessary, adaptive component of social behavior. In humans, however, some forms of aggression are considered pathological when they threaten lives, increase the risk of psychiatric impairment in victims pointing to the need for identifying targets for therapeutic interventions (Golden, Jin, Heins, et al., 2019; Golden, Jin, & Shaham, 2019; Kessler et al., 2006; McCloskey et al., 2010). Research on aggression has focused on a few brain regions termed the “core aggression circuit” (CAC) in mostly male mice (Mei et al., 2023). The CAC largely overlaps with the “social behavior network” which was largely identified by examining c-Fos in a handful of regions that expression steroid hormone receptors (Goodson, 2005; Lawal et al., 2025; Newman, 1999) such as the medial amygdala (Hong et al., 2014; Unger et al., 2015), the ventrolateral portion of the ventromedial hypothalamus (Dai et al., 2025; Falkner et al., 2016; Hashikawa et al., 2017; Lee et al., 2014; Lin et al., 2011; Mountoufaris et al., 2024; Yan et al., 2024), the posterior portion of the bed nucleus of the stria terminalis (Bayless et al., 2019; Yang et al., 2022), and the ventral pre-mammillary nucleus (Chen et al., 2020; Stagkourakis et al., 2018). Recent work also indicates that regions of the pallial amygdala such as the posterior amygdala (Yamaguchi et al., 2020; Zha et al., 2020) and the posterolateral cortical amygdala (Aubry et al., 2025) are also part of this network.

While studying these regions has led to great progress in understanding the neural circuits of aggression, the mouse brain contains hundreds of regions, which leaves open the possibility that the network of regions which play a significant role in aggression is larger than expected (Simpson et al., 2021). Unbiased, whole-brain approaches are therefore needed to reveal how large-scale functional networks reorganize depending on the behavior the animal engages in. Indeed, recent c-Fos mapping combined with network analyses has demonstrated how interregional coactivation patterns are altered as a result of fear memory consolidation (Roy et al., 2022; Vetere et al., 2017) and exposure to alcohol (Ardinger et al., 2024; Kimbrough et al., 2020; Roland et al., 2023), complementing region-focused analyses and reducing a priori bias toward canonical circuits.

In this paper, we used whole-brain c-Fos mapping combined with a behavioral paradigm which elicits aggression in both male and female outbred Swiss Webster mice (Aubry et al., 2022; Newman et al., 2019). We adopted the Weighted Gene Co-expression Network Analysis (WGCNA) framework (Langfelder & Horvath, 2008; Zhang & Horvath, 2005) to our c-Fos data to identify mesoscale modules of co-activation in aggressive (AGG) and non-aggressive (NON) males and females. Module preservation analysis was performed between AGGs and NONs within each sex using permutation-based preservation statistics of module density and connectivity to test which modules were reorganized in AGGs relative to NONs (Langfelder et al., 2011). Finally, within modules that show altered organization, we perform differential correlation to localize the specific region pairs whose adjacencies differ significantly between AGG and NON (McKenzie et al., 2016).

## Results

### Social Behavior in male and females

We quantified various social behaviors on the day mice were euthanized for whole brain quantification of c-Fos expression (Fig. 1A). As expected, we found that both AGG male and female mice engaged in aggressive behavior for a longer period of time than NONs (F_(1,69)_ = 54.14, p < 0.0001, Fig 1B). When examining gross interaction, we found that females (F_(1,69)_ = 27.62, p < 0.001) engaged in more investigation than males of each phenotype (NONs: p < 0.001, AGGs: p = 0.0315, Fig. 1C). We next examined investigation of distinct body parts and found that females investigated the intruders anogenital region (F_(1,69)_ = 14.87, p = 0.0003), face (F_(1,69)_ = 33.63, p < 0.0001) and flank (F_(1,69)_ = 7.330, p = 0.0085) more than males. We also found a significant Phenotype x Sex interaction for facial investigation (F_(1,69)_ = 6.591, p = 0.014) and flank investigation (F_(1,69)_ = 5.346, p = 0.0238). Post-hoc comparisons revealed that female AGGs engaged in more facial investigation than female NONs (p = 0.0459) and male AGGs (p < 0.0001). In contrast, female NONs engaged in more flank investigation than female AGGs (p = 0.0058) and male NONs (p = 0.0019). When examining distinct attack behaviors, we did not find a difference between biting (t_(33.91)_ = 0.04829, p = 0.9618) or kicking (t_(29.46)_ = 1.829, p = 0.0776), but males engaged in more wrestling behavior than females (t_(21)_ = 2.316, p = 0.0307).

**Figure 1.**
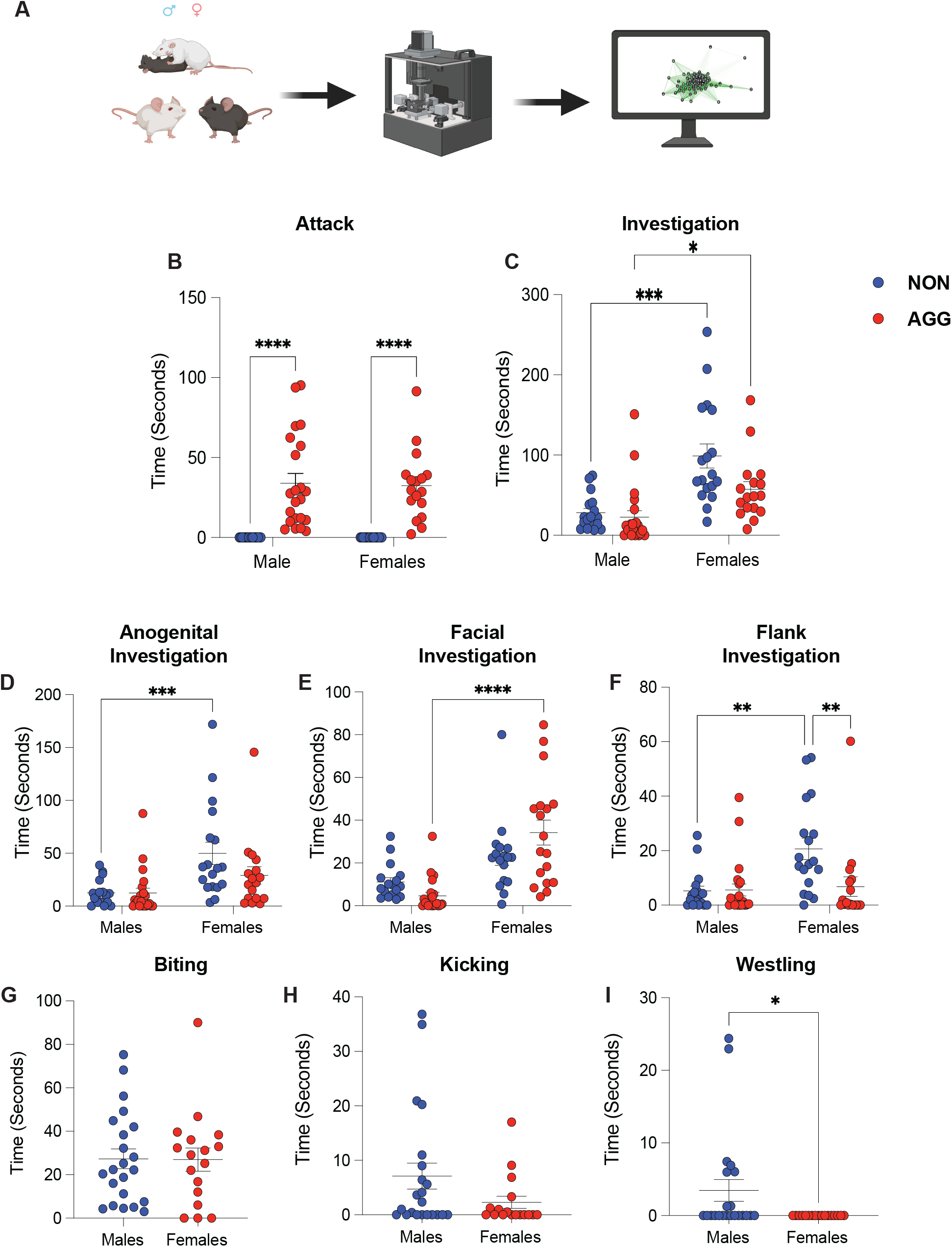
Social behavior prior to whole-brain c-Fos quantification. (A) Timeline: resident–intruder assay followed by perfusion and iDISCO+ whole-brain c-Fos imaging. (B) Total time spent in aggressive behavior: AGG mice (male and female) engaged in aggression longer than NONs. (C) Gross social investigation: females investigated more than males across phenotypes (D–F) Investigation by body region: females investigated the intruder’s (D) anogenital region and (F) flank more than males. (G–I) Attack subtypes: no sex differences in (G) biting or (H) kicking but (I) males wrestled more than females Data are shown as group means ± SEM with individual data points; two-way ANOVA with post-hoc comparisons for panels B–F, unpaired t-tests for panels G–I. Significance defined at p<0.05. n = 17 Female AGG, 17 Female NON, 22 Male AGG, 17 Male NON

### Differential c-Fos Expression

We first examined differential c-Fos expression between the two phenotypes within males and females. Overall (Fig 2A), males AGGs showed a higher number of regions with an increase in c-Fos compared to NONs (76, 29%) than female AGGs (16, 6%). When examining specific points in space along the anterior-posterior (AP) axis we observed striking differences between male and female AGGs relative to the NONs.

**Figure 2.**
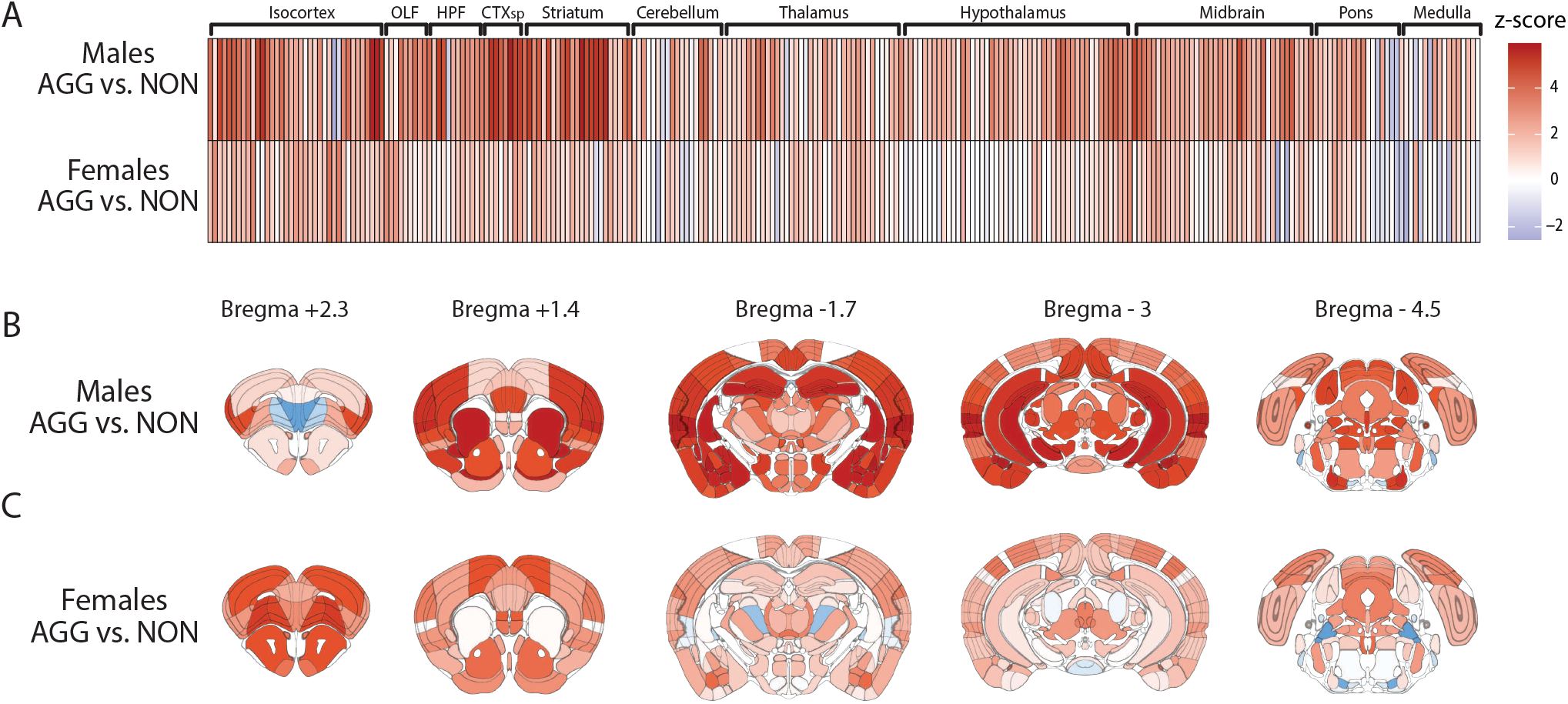
Phenotypic-specific patterns of aggression-evoked c-Fos activation. (A) Heatmap displaying z-scored differences between male AGGs and NONs (top) and female AGGs and NONs (bottom). (B) Z-score overlayed on to 5 different coronal sections across the anteroposterior axis comparing male AGGs and NONs. (C) ) Z-score overlayed on to 5 different coronal sections across the anteroposterior axis comparing female AGGs and NONs n = 22 Male AGG, 17 Female AGG, 17 Male NON, 17 Female NON

Relative to male NONs, AGGs showed an increase in brain regions throughout the AP axis (Fig 2B). The largest increases were observed in regions the amygdala complex (Bregma – 1.7) including all sub-regions of the basolateral amygdala, the cortical amygdala, the posterior amygdala and medial amygdala (Bregma – 1.7). The insular, temporal association, endopiriform, and ectorhinal cortices also showed large differences in c-Fos expression (Bregma + 2.3, 1.4, -1.7, -3). Other notable increases were found in hypothalamic areas such as the ventromedial hypothalamus (Bregma -1.7) and midbrain areas including the superior colliculus, substansia nigra and ventral tegmental area (Bregma -3, -4.5). In contrast, almost all of the regions that showed an increase in c-Fos in female AGGs relative to NONs (Fig. 2C) were found to be in anterior cortical regions including the infralimbic, orbital and supplementary motor cortices in addition to anterior olfactory regions such as the dorsal peduncular, anterior olfactory nucleus and the taenia tecta (Bregma + 2.3). Female AGGs also showed an increase in c-Fos in the nucleus accumbens (Bregma +1.4), the ventral basolateral amygdala (Bregma -1.7) and in the deep layers of the superior colliculus (Bregma -4.5). For a full list of regions see Supplementary Table 1.

We next compared differences in c-Fos expression between the sexes within each phenotype (Fig. 3A). When comparing AGGs, we found that males showed an increase in c-Fos in 78 total regions (29%) and females showed an increase in 7 regions (3%). When comparing NONs, we found that males showed an increase in 19 regions (7%) and females showed an increase in 8 (3%). As with the comparison to male NONs, male AGGs showed increases in c-Fos relative to females primarily in the amygdala complex with particularly strong increases in the lateral, posterior basolateral, and the cortical amygdala (Bregma -1.7). Males also showed strong increases in all areas of the hippocampal formation (Bregman -1.7, -3), the ventral tegmental area (Bregma -3), superficial layers of the superior colliculus and the inferior colliculus (Bregma -4.5). Regions in which females showed greater c-Fos expression than males were concentrated in anterior cortical regions (Fig. 3B). These regions included all three sub-regions of the orbital cortex, the prelimbic, infralimbic and supplementary motor cortices (Bregma +2.3). Within the NONs, males showed an increase in c-Fos primarily in the retrosplenial cortex and numerous areas of the visual cortices (Bregma -3, -4.5). Female NONs showed increases in the primary motor and anterior cingulate cortices as well as the caudate putamen and the endopiriform cortex. (Bregma +1.4). For a full list of regions see Supplementary Table 2.

**Figure 3.**
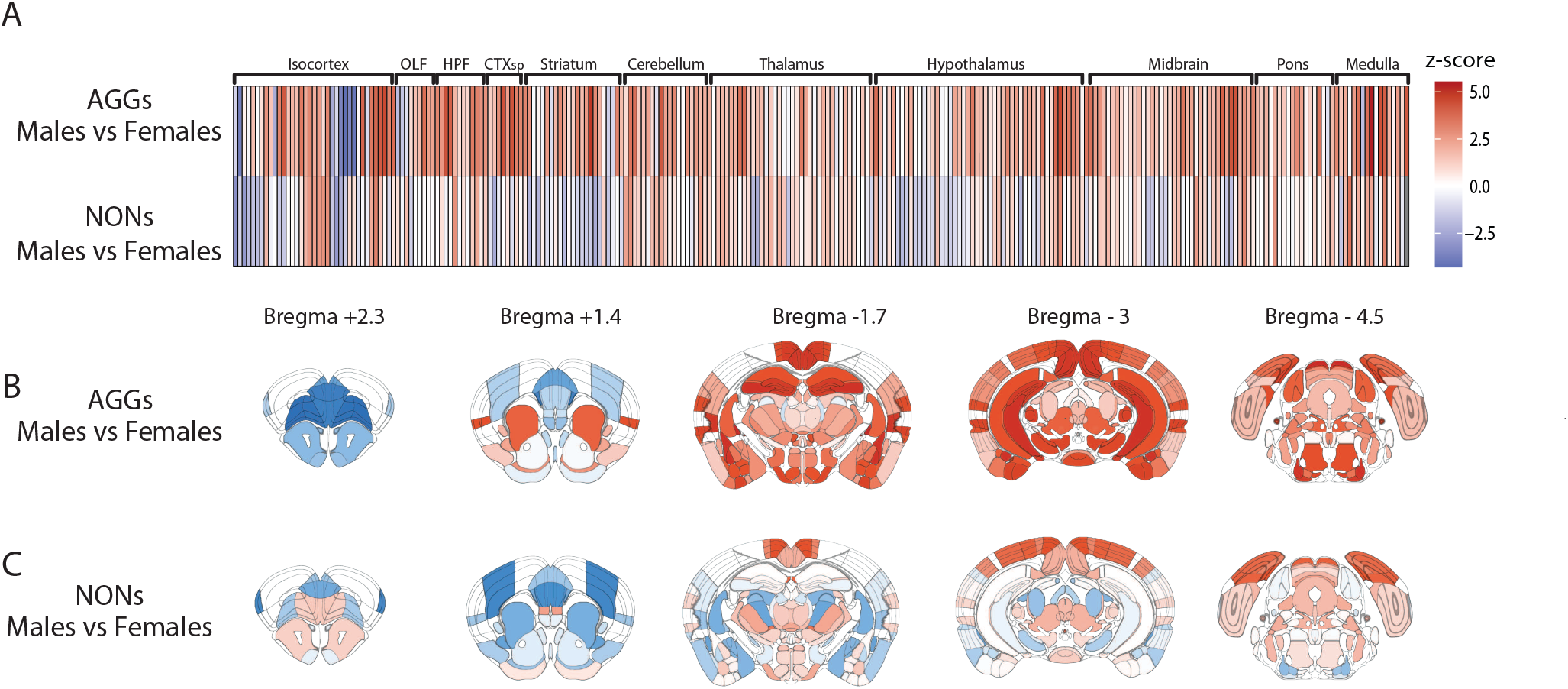
Sex-specific patterns of aggression-evoked c-Fos activation. (A) Heatmap displaying z-scored differences between male and female AGGs (top) and NONs (bottom). (B) Z-score overlayed on to 5 different coronal sections across the anteroposterior axis comparing male and female AGGs. (C) Z-score overlayed on to 5 different coronal sections across the anteroposterior axis comparing male and female NONs. n = 22 Male AGG, 17 Female AGG, 17 Male NON, 17 Female NON \

### Whole Brain Network Analyses

To understand how these regions are co-activated in male and female AGGs, we applied the weighted-gene co-expression network analysis (WGCNA) framework to our data set. We constructed a network for each of the four groups and then preformed a module preservation analysis for each sex with the AGG network set as the reference network. We found that the NON networks for each sex were able to preserve the density of connections in the identified modules, however we found that the specific connectivity profiles were not preserved in select modules from male AGGs.

In males, the blue module (Fig. 4A) was a large cluster linking together numerous cortical areas with the basal ganglia, septum, thalamic relay areas and superior colliculus sub laminae. Of the 77 total regions in this cluster, 66 showed significant differences in their total connection strength with other regions in the module. Of these 66, AGG mice showed higher connectivity strengths in 63 of the 66 (Fig. 4B). Given that the connectivity profiles include both direct connections between each pair of regions and their shared connections with all other nodes in the cluster, we performed a differential correlation analysis on the regions which showed significantly different connection strengths to examine pairwise correlations. Of the 2016 correlations evaluated, 309 showed a significant difference between the AGG and NON modules (Fig. 4C). Of the 309 significantly different correlations, 299 were stronger in AGGs than NONs. The yellow module (Fig. 4D) spanned olfactory cortical areas and brainstem regions. Of the 40 regions, 30 showed a significant difference in total connectivity, all of which were higher in the AGG module (Fig. 4E). Of the 435 correlations examined, 177 were significantly different between AGGs and NONs, 173 of which showed higher correlations in AGGs (Fig. 4F).

**Figure 4.**
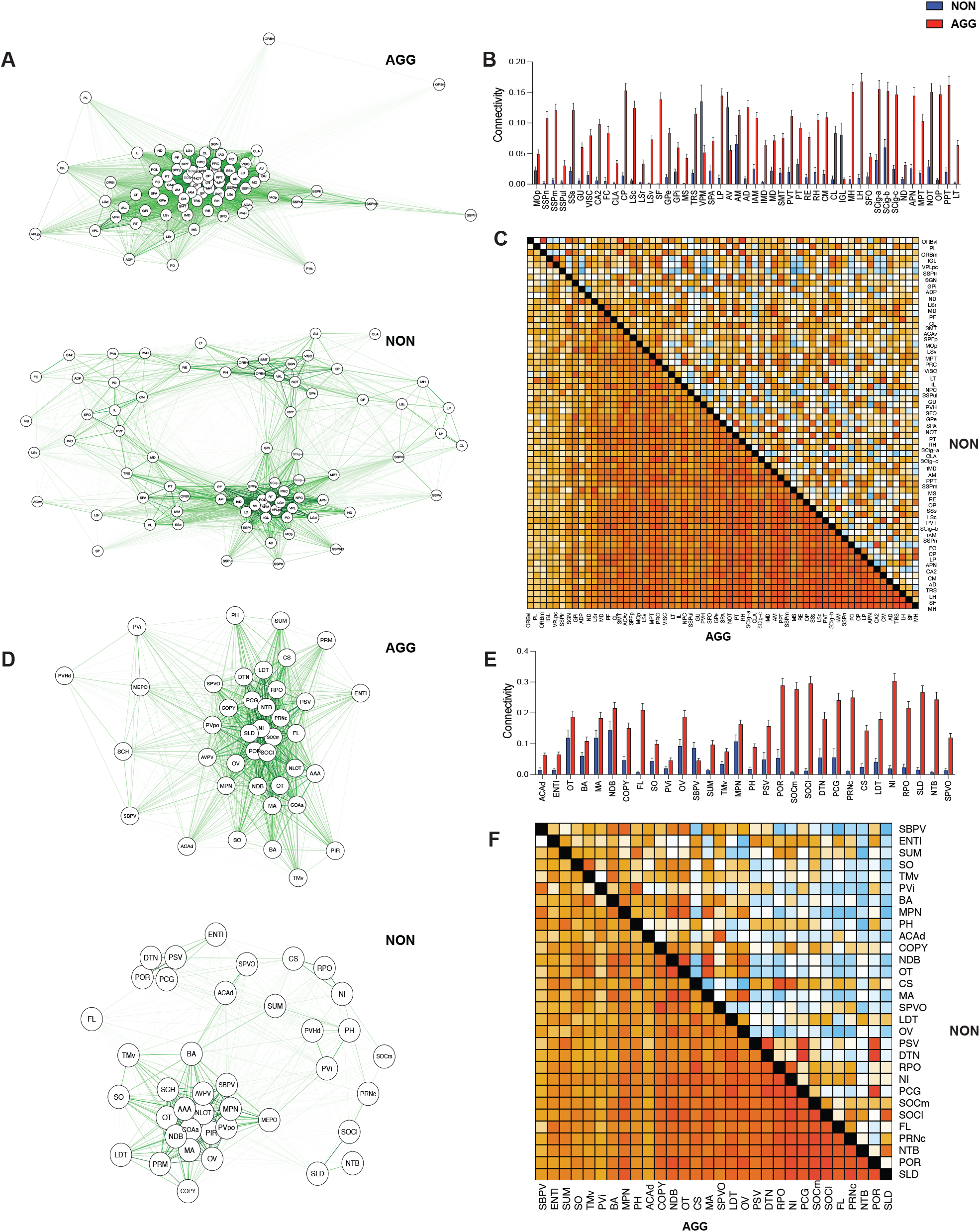
The blue and yellow modules were differentially connected in male AGGs vs. NONs. (A) Network plot of the blue module in AGGs (top) and NONs (bottom). (B) Comparison of total intramodule connectivity in AGGs and NONs for the blue module. (C) Edge-wise differential correlation analysis within the blue module in AGG vs NONs. Warmer colors indicate positive correlations; cooler colors indicate negative correlations. Heatmap below the diagonal represents correlations in the AGGs, heatmap above the diagonal represents correlations in the NONs. (D) Network plot of the yellow module in AGGs (top) and NONs (bottom). (E) Comparison of total intramodule connectivity in AGGs and NONs for the yellow module (F) Differential correlation analysis within the yellow module AGG vs NONs.

The green network was a brainstem-heavy sensorimotor network with a few multimodal cortical regions. Of the 36 regions, 25 regions showed an increase in connectivity in the AGG module (Fig. 5A), with AGGs showing increased connectivity in 23 of the 25 (Fig. 5B). When examining the individual correlations in the module 81 of 300 were significantly different between groups, 76 of which were higher in AGGs (Fig. 5C). Although the connectivity profile was moderately preserved, we included the brown module (Fig. 5D) in our analysis as it includes many of the regions which have been identified as the “social behavior network”, such as the VMH, BNST, MEA, COAp, PMv, AHN and the MPOA. We found that AGGs had a significant increase in total connectivity in 35 of the 45 regions (Fig.5E). When examining the individual correlations, we found that 89 of the 630 correlations examined were significantly different, all of which were higher in AGGs (Fig.5F).

**Figure 5.**
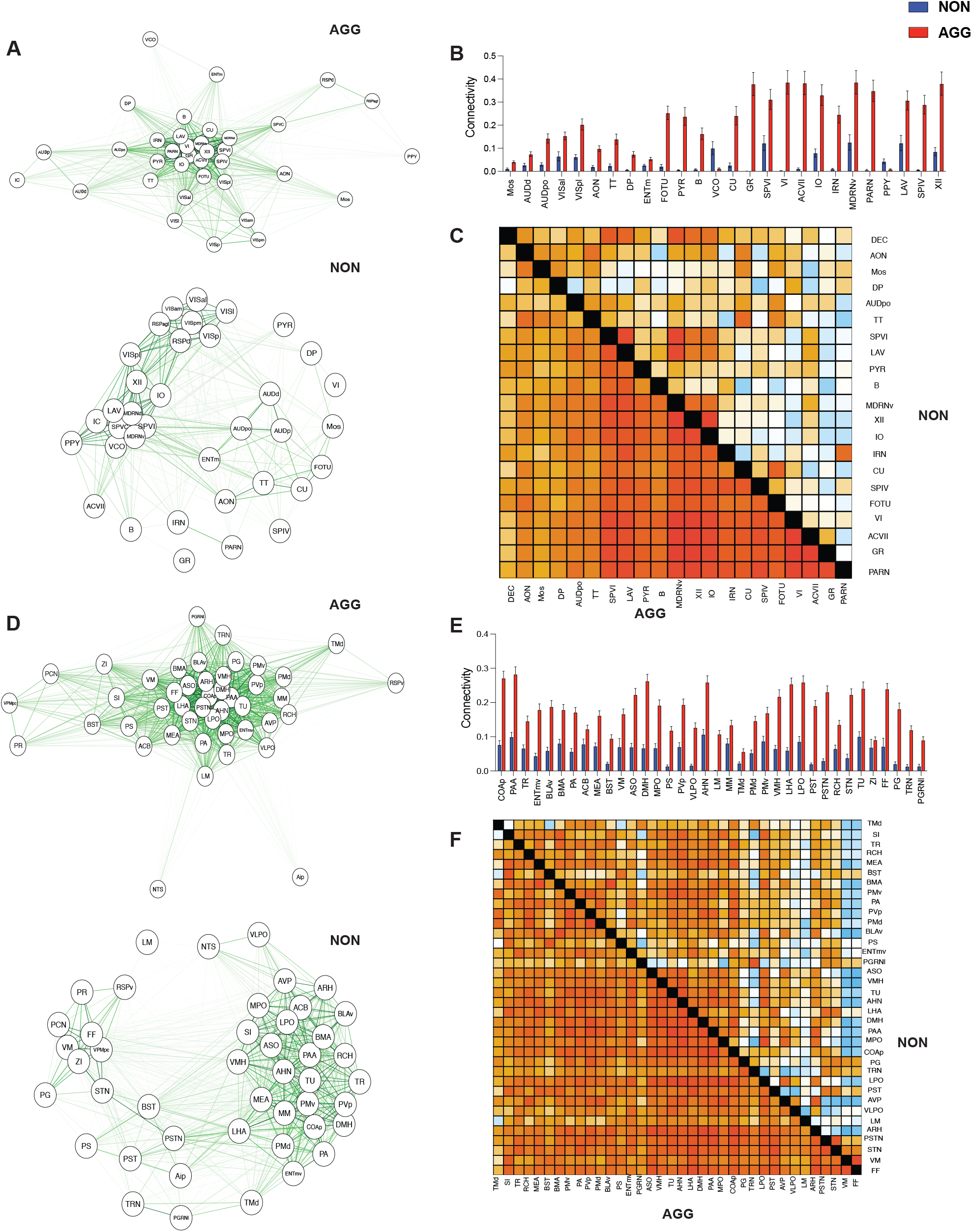
Aggression strengthens connectivity in the green and brown modules in males. (A) Network plot of the green module in AGGs (top) and NONs (bottom). (B) Comparison of total intramodule connectivity in AGGs and NONs for the green module. (C) Edge-wise differential correlation analysis within the green module in AGG vs NONs. Warmer colors indicate positive correlations; cooler colors indicate negative correlations. Heatmap below the diagonal represents correlations in the AGGs, heatmap above the diagonal represents correlations in the NONs. (D) Network plot of the brown module in AGGs (top) and NONs (bottom). (E) Comparison of total intramodule connectivity in AGGs and NONs for the brown module (F) Differential correlation analysis within the brown module AGG vs NONs

When examining the female networks, we found that all AGG modules were moderately preserved in NONs based on the z-scored based metrics derived from permutation testing. Therefore, we chose to analyze the two modules with the weakest connectivity preservation based on the observed metrics. The turquoise module spanned somatosensory/interoceptive cortices, hippocampus, thalamus, and multiple regions in the midbrain. Of the 55 regions in the module 30 showed significant differences in total connectivity, 24 of which were increased in AGGs (Fig 6A-6B). Of the 465 correlations analyzed, 56 were significant with 47 showing higher correlations in AGGs. The red module contained many of the frontal cortical regions identified in the differential expression analysis as well as striatal and thalamic regions. Of the 28 regions in this module, only 5 showed significant differences in total connectivity strength, all of which were increased in AGGs. Given the low number of regions with differences in connectivity strength, we analyzed all possible pairwise correlations and found that only 3/379 region pairs showed differential correlations, all of which were increased in female AGGs.

**Figure 6.**
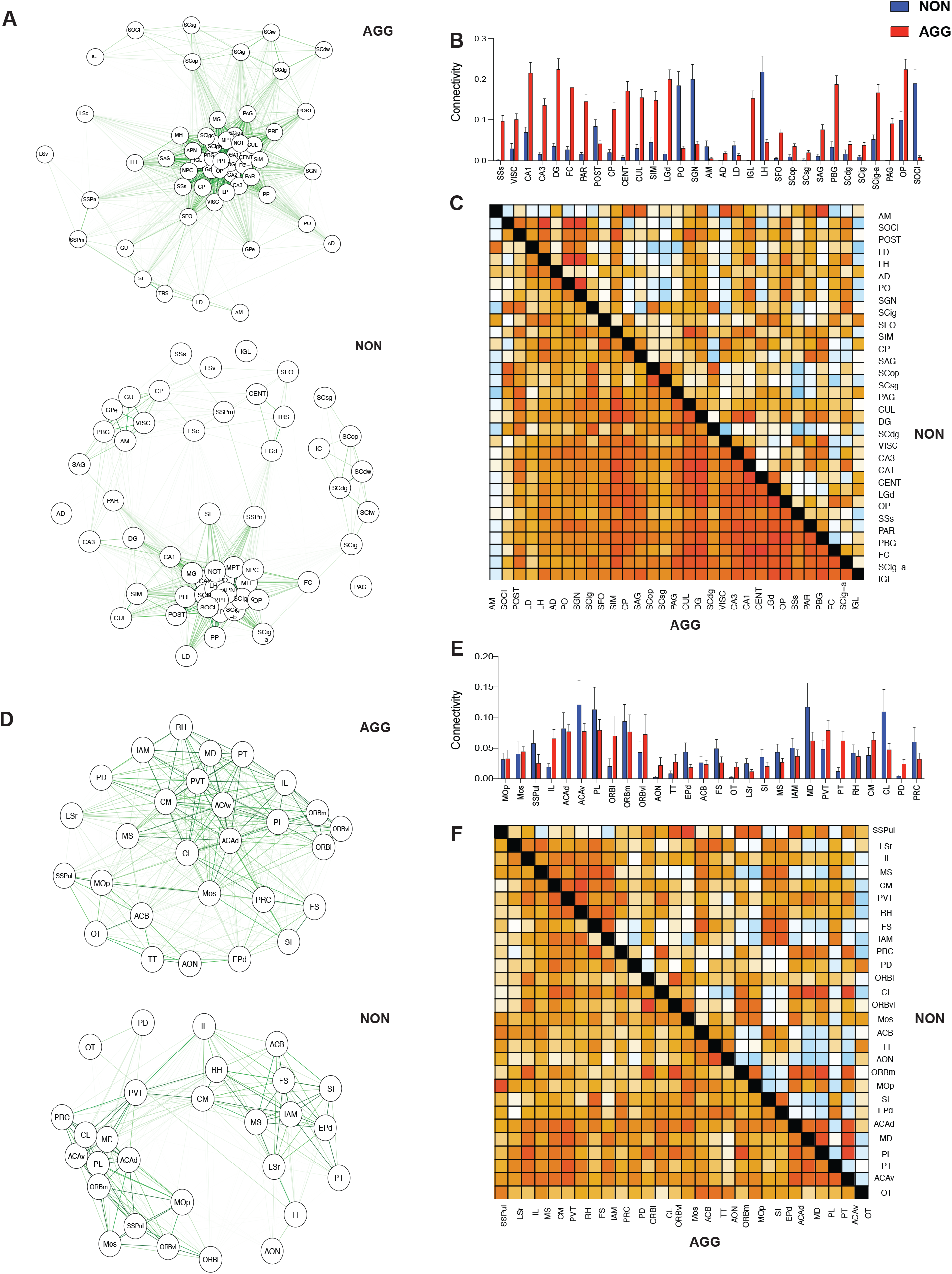
Turquoise and red modules were the least preserved modules in females. (A) Network plot of the turquoise module in AGGs (top) and NONs (bottom). (B) Comparison of total intramodule connectivity in AGGs and NONs for the turquoise module. (C) Edge-wise differential correlation analysis within the turquoise module in AGG vs NONs. Warmer colors indicate positive correlations; cooler colors indicate negative correlations. Heatmap below the diagonal represents correlations in the AGGs, heatmap above the diagonal represents correlations in the NONs. (D) Network plot of the red module in AGGs (top) and NONs (bottom). (E) Comparison of total intramodule connectivity in AGGs and NONs for the red module (F) Differential correlation analysis within the red module AGG vs NONs

## Discussion

This study reveals that aggressive social encounters reorganize brain-wide c-Fos activation in a sex-dependent manner. Despite similar levels of attack duration in both males and females, males showed a greater of c-Fos activity than females. Males displayed an increase in c-Fos in 78 regions (29%) compared to female AGGs and in 76 (29%) regions compared to male NONs. In contrast, females only exhibited increases in c-Fos in 7 (3%) regions compared to male AGGs and 16 (6%) compared to female NONs. Specifically, major sex differences were observed in the cortex and amygdala, with male AGGs showing significant increases in the somatosensory, interoceptive, and posterior polymodal cortices, while females showed preferential activation of anterior cortical regions. In addition, males demonstrated a broad increase across the entire amygdala complex with females only activating the BLAv and BMA. There were also striking differences in the olfactory regions activated in male and female AGGs. Males activated olfactory regions such as the piriform cortex, cortical and medial amygdala, and the pirifrom-amygdala area while females activated the anterior olfactory nucleus, the dorsal peduncular area and the taenia tecta. Together, these results suggest that aggression induces a reconfiguration of distributed mesoscale structures whose composition differ by sex. Males show a broad level of activation throughout the AP axis while females primarily activate anterior olfactory and cortical areas.

There are two possible explanations for the differences in total number of regions activated in males results at the behavioral level. First, although there were similar levels quantitatively, the patterns of aggression are qualitatively different. Males display explosive bouts of attack which include wrestling and lunging towards the intruder. Males also find these bouts reinforcing as evidenced by forming a conditioned place preference and lever pressing for the chance to engage in fighting while female mice do not (Aubry et al., 2022). It is possible that in a different social context, such as maternal aggression, we would see higher levels of c-Fos activation throughout the brain in females given the evolutionary importance of this type of aggression. Evidence for this explanation comes from previous work identifying the cortical, posterior, and medial amygdala, medial preoptic nucleus, and the ventromedial hypothalamus as regions activated by maternal aggression (Gammie & Nelson, 2001; Yamaguchi et al., 2026), regions which did not show significant activation compared to female NONs in our paradigm. We chose not to use maternal aggression in our study since this model is not ideal for evaluating sex differences given that these behaviors are linked to hormonal changes specifically associated with pregnancy, parturition, and lactation. Future studies are thus needed to determine how maternal aggression differs from the rival aggression model used in this study (Newman et al., 2019).

We also constructed separate networks for AGG and NON males and females, then tested whether AGG-defined modules were preserved in NON modules within each sex. Methodologically, WGCNA offered clear advantages over simple correlation maps. First, soft-thresholded, signed adjacency and topological overlap measures down-weight spurious connections and accentuate shared-neighborhood structure, yielding more stable modules than binarized or thresholded matrices (Zhang & Horvath, 2005). Second, permutation-based Z statistics disentangle density-level from connectivity-level preservation, highlighting modules that look intact in aggregate yet hide substantial rewiring of which region occupies central positions (Langfelder et al., 2011). Preservation analysis indicated that NON networks maintained the overall density of several AGG modules but failed to preserve the specific intra-module connectivity profiles in select cases suggesting that aggression reorganizes the connectivity profiles even when aggregate density is similar. Density-level preservation paired with connectivity-level changes suggest large-scale rewiring rather than simple up- or down-scaling of the same edges.

In males, the blue module exemplified large-scale strengthening with aggression. This module linked sensorimotor and interoceptive cortices with frontal areas, basal ganglia, septum, thalamic relays, and the superior colliculus. A large majority of its regions showed altered total connectivity, and most of those were stronger in AGG than NON, indicating module-wide connectivity increases. The yellow module (olfactory cortex, pallidal, preoptic/hypothalamic, and brainstem nodes) showed a similar pattern with three quarters of regions increased total connectivity in AGG and most significant edges were stronger with aggression—again consistent with a high-gain state. The green, brainstem-heavy sensorimotor module also strengthened in AGG at both the region and edge levels. Surprisingly, the connectivity profile brown module which was composed of classic social-behavior regions (VMH, BNST, MEA, COAp, PMv, AHN, MPOA, AVPV) was moderately preserved. That there were increases in total connectivity, with relatively small differences in region specific correlations suggests that this module may simply scale-up activity in AGGs relative to NONs with changing functional connectivity patterns. This is in contrast to the other modules which showed broad changes in both total connectivity and the re-routing of paired correlations

By contrast, female AGG networks were conserved to a greater degree than males. Preservation statistics indicated that AGG modules were generally maintained in NONs, prompting focused analyses on the two least-preserved modules. The turquoise module (somatosensory/interoceptive cortex, hippocampus, lateral septum, thalamo-visual relays, multiple SC layers, and midbrain regions) showed selective increases in about half of the and most significant edges were higher in AGGs. Interestingly, the red module consisted of mainly of regions in the anterior portions of the cortex such as the PL, IL, OFC and the DP and AON which are located ventral of the PL/IL. All of these regions also showed increases in c-Fos counts relative to male AGGs and female NONs. This finding is particularly interesting in light of previous optogenetic and lesion studies showing that the mPFC inhibits aggression in male rodents (de Bruin et al., 1983; Kolb & Nonneman, 1974; Takahashi et al., 2014). This is perhaps the most striking sex difference observed in this study and warrants further in-depth study in the future.

There are limitations in this work which warrant consideration. First, c-Fos is an indirect proxy of recent neuronal activation, lacks the temporal resolution, the ability to determine cell-type specific activation, and is unable to detect inhibition (Kovacs, 2008; Yang et al., 2024). These caveats point directly to future work. Causal tests in identified hubs could assess whether boosting or suppressing their activity shifts module structure and behavior in a sex-specific manner. Additionally, multisite recordings with either photometry or electrophysiology would go a long way in confirming the accuracy of the modular structures identified by our network analyses (Guo et al., 2023; Mague et al., 2022). Additionally, integrating structural connectivity could help distinguish direct from polysynaptic co-activation.

In summary aggression recruits distributed mesoscale networks, and this recruitment differs by sex. By leveraging WGCNA with module preservation and differential correlation, we uncovered modules that are rewired in their internal connectivity, and we pinpoint the edges most sensitive to aggressive state. These insights provide a system-level framework for selecting candidate hubs and subnetworks for causal interrogation and, ultimately, for understanding how the brain orchestrates complex social behavior in a sex-specific manner.

## Methods

### Animals

Wild-type Swiss-Webster (SW) mice (male and female, 12-15 weeks; Charles River) were used as experimental (residents) mice. Intruders for the resident-intruder (RI) test were 8-12-week-old male or female C57/BL6J mice (Jackson Laboratory). All mice were allowed to acclimate to the housing facility for 1 week prior to any experimental protocol. Mice were paired with a member of the opposite sex for sexual experience at 9 weeks of age (females) for two weeks or at 11 weeks of age (males) for two days (Aubry et al., 2022). Females were paired with castrated SW males (8-12 weeks; Charles River) to prevent pregnancy (Newman et al., 2019). All mice were maintained on a 12/12 h light/dark cycle (07:00–19:00) with *ad libitum* access to food and water. Housing and experimental rooms were maintained at 20–22 °C and 40–60% humidity. Experiments were conducted during the light phase. Procedures were performed in accordance with the National Institutes of Health Guide for Care and approved by the Use of Laboratory Animals and the Icahn School of Medicine at Mount Sinai Institutional Animal Care and Use Committee.

### RI Test

Mice were screened using protocols adapted from previous studies(Aubry et al., 2022; Li et al., 2023). Cage tops were removed to monitor the final trial with an overhead camera (Sony HDR-CX440). A novel C57BL/6J mouse matching the sex of the resident was introduced into each cage and mice were allowed to freely interact for 5 min across three days. After 5 min elapsed, intruder mice were returned to their home cages. For female resident-intruder trials, cohabiting male mice were removed prior to the test and then returned to their home cages. Resident behaviors were manually annotated using Observer XT 11.5 (Noldus Interactive Technologies).

### Perfusion, brain tissue processing, and imaging

For iDISCO+, mice were injected with 10% chloral hydrate and perfused transcardially with ice-cold 1× PBS (pH 7.4) following the third RI test. Tissue was fixed with cold 4% paraformaldehyde in 1× PBS. Brains were post-fixed for 12 h in the same fixative at 4 °C and then transferred to 1X PBS. The iDISCO+ staining protocol was modified from http://www.idisco.info. Fixed whole brains were incubated with the primary c-Fos antibody (no. 226003, 1:1,000, Synaptic Systems) and secondary donkey anti-rabbit IgG (H+L) Highly Cross-Adsorbed Secondary Antibody, Alexa Fluor 647 (no. A-31573, 1:1,000, Thermo Fisher Scientific) for 7 days each. A LaVision light sheet microscope with zoom body was used for half-brain sagittal scanning, with dynamic focus and a step size of 4 μm.

### Regional Count Analysis

Cleared brains were processed as previously described using Clear Map 1 (Renier et al., 2016; Renier et al., 2014). c-Fos^+^ cells were quantified using the cell detection module, with cell detection parameters optimized and validated based on the intensity and shape parameters of the signal. The autofluorescence channel was aligned to the Allen Institute’s Common Coordinate Framework using the Elastix toolbox. We analyzed the resulting counts with a negative binomial generalized linear model using the MASS package in R. For each fitted model, we obtained estimated marginal means and performed pairwise contrasts for sex and phenotype. We controlled for FDR using the Benjamini and Hochberg procedure in Prism (version 10.6.0). Significance was set at q <0.10. Heatmaps demonstrating phenotypic and sex differences were generated using the napari and brainglobeatlas packages in python.

### Network and module preservation analysis

To identify c-Fos co-expression modules we performed Weighted Gene Co-expression Network Analysis (WGCNA) using the WGCNA R package (Langfelder and Horvath, 2008). A signed weighted adjacency matrix was constructed by computing pairwise Pearson correlations between all region pairs and raising the absolute value of each correlation to a soft-thresholding power β to approximate scale-free topology. The optimal β was selected using the pickSoftThreshold function to ensure that each network exhibited approximate scale-free topology (R^2^ > 0.8). The adjacency matrix was transformed into a Topological Overlap Matrix (TOM), which measures the overlap in shared network neighbors between regions and reduces the influence of spurious or isolated connections. Regions were hierarchically clustered using average linkage on 1 – TOM dissimilarity, and modules were identified via dynamic tree cutting at a height of .25. Each module was assigned a unique color label and summarized by its module eigen-region, defined as the first principal component of the scaled expression matrix for regions within the module.

To assess differential connectivity of sex specific AGG and NON networks and their stability within datasets, we conducted module preservation analysis using the modulePreservation function. This analysis evaluates whether modules defined in a reference dataset (AGG) are preserved in a test dataset (NON), based on both network connectivity and density statistics. It also allows for the assessment of cluster quality based on the consistency of module structure across randomly partitioned subsets of the reference data. To assess the quality and preservation of modules, Z statistics were computed for each module. Each Z-score was calculated by comparing the observed module statistics to a null distribution generated from 500 permutation tests where module labels or sample labels were randomized. For a given statistic, the Z-score is calculated as: Z = ((observed – μ_null) / σ_null) where μ_null and σ_null are the mean and standard deviation of the statistic under the null distribution. Larger Z-values indicate greater evidence of preservation or quality. The four z-statistics which comprises the density metric are:

meanSignAwareCor: measures the mean pair-wise signed Pearson correlations among every region pair inside the module in the test set. Strong preservation of this statistic indicates that the overall correlation (strength and direction) of the module remains tighter than assumed by the null distribution.

meanAdj: the average edge weight among module regions in the test network. A large Z for this statistic shows that the module still forms a densely connected block compared with modules generated by label permutations.

propVarExplained: quantifies how much of the within-module expression variance in the test data is still captured by the module eigen-region. A high, positive Z-score means the regions continue to behave collectively along a dominant expression axis.

meanSignAwareKME: evaluates the average signed module-membership (kME) values of all module genes in the test network. By retaining the sign information of each gene’s correlation with the module eigengene, it asks whether genes remain positively or negatively aligned with that eigengene more strongly than in permuted data.

The z-statistics which comprise the connectivity measure are:

Zcor.KIM: evaluates the Pearson correlation between intramodular connectivity (kIM) vectors calculated in the reference and test networks. For each region, kIM is the sum of its edge weights to all other regions in the module. A high, positive Z-score indicates that the rank order of hubs versus peripheral genes is preserved far better than would be expected under random label permutations.

ZcorKME: captures the correlation between module-membership scores (kME) across the two datasets. kME is each region’s correlation with the module eigen-region which reflects how strongly a region’s expression follows the module’s dominant expression pattern. A high Z for this statistic means regions that were eigen-region aligned in the reference set remain similarly aligned in the test set, preserving the expression-centric notion of centrality.

Zcor.cor: measures the correlation between the entire matrices of pair-wise gene-gene correlations within the module in the reference and test data. Conceptually, all within-module correlations are flattened into a single vector for each dataset, and those two vectors are then correlated. A large Zcor.cor value signals that the strength of the correlations between regions in the module are preserved beyond chance.

We tested edge-level differences in functional coupling between aggressive and non-aggressive animals using Differential Correlation analysis with the DGCA framework (McKenzie et al., 2016). For each group, we assembled region × animal matrices of c-Fos activity and computed Pearson’s correlation for every region pair. Correlations were z transformed, and edges were compared between groups using DGCA’s two-sample z-test. Multiple testing was controlled with Benjamini–Hochberg FDR; significant edges (q < 0.10).

### Statistical analysis

All statistical tests and associated information are reported in the Figure legends. All *t*-tests, one-two-way ANOVAs were performed using GraphPad Prism software (GraphPad Software Inc. version 10.6.0). Two-way ANOVA analysis was followed by Šídák’s multiple-comparisons test for post-hoc analysis. For comparisons between groups for region by region iDISCO+ analysis, *P* values were corrected for multiple comparisons using the Benjamini–Hochberg procedure to reduce false discovery rate. *Q* values below 0.10 were considered significant.

## Acknowledgements

We thank A. Smith for help with the iDISCO+ protocol and Clear Map analysis, and N. Tzavaras, K. Cialowicz and G. Doherty for help with light-sheet imaging. This work was supported by the US National Institutes of Health (grants R01MH114882, R01MH127820, R01MH104559, R01MH120514 and R01MH120637 to S.J.R K99DA058213 to A.V.A.); the Doft family for the Friedman Brain Institutes’ Postdoc Innovator award (A.V.A.); the Brain and Behavior Research Foundation (grants 30233 to L.L., 31140 to R.D.-d.C., 30894); the Beijing Natural Science Foundation (grant Z240012 to L.L; and the NIH Virus Center (grant P40 OD010996).

## Author Contributions

Conceptualization: A.V.A and S.J.R. Investigation: A.V.A, L.L, R.D.C, E.K, H.O, A.T. Data Analysis: A.V.A, Writing-original draft: A.V.A, reviewing and editing: A.V.A, R.D.C, L.L, S.J.R. Supervision: A.V.A and S.J.R

